# Mycobacterial OtsA structures unveil substrate preference mechanism and allosteric regulation by 2-oxoglutarate and 2-phosphoglycerate

**DOI:** 10.1101/677302

**Authors:** Vítor Mendes, Marta Acebrón-García-de-Eulate, Nupur Verma, Michal Blaszczyk, Márcio V. B. Dias, Tom L. Blundell

## Abstract

Trehalose is an essential disaccharide for mycobacteria and a key constituent of several cell wall glycolipids with fundamental roles in pathogenesis. Mycobacteria possess two pathways for trehalose biosynthesis. However, only the OtsAB pathway was found to be essential in *M. tuberculosis*, with marked growth and virulence defects of OtsA mutants and strict essentiality of OtsB2. Herein, we report the first mycobacterial OtsA structures from *M. thermoresistibile* in both apo and ligand-bound forms. Structural information reveals three key residues in the mechanism of substrate preference that were further confirmed by site-directed mutagenesis. Additionally, we identify 2-oxoglutarate and 2-phosphoglycerate as allosteric regulators of OtsA. The structural analysis in this work strongly contributed to define the mechanisms for feedback inhibition, show different conformational states of the enzyme and map a new allosteric site.

## Introduction

Trehalose is a non-reducing disaccharide, formed by α-(1-1) linked glucoses, with a wide distribution in nature and present in all three domains of life (1). This remarkably widespread sugar performs multiple roles in a wide variety of organisms and it can also fulfil different roles within the same organism. Trehalose has been considered a compatible solute, conferring protection to proteins, DNA, membranes and whole cells from thermal shock, osmotic shock, freezing, ionizing radiation, oxidative stress and desiccation (1-9). This disaccharide, which can further function as a carbon and energy reserve molecule (1, 10-12), was also recently related to pathogenicity of *Pseudomonas aeruginosa* in plants (13). It additionally plays fundamental signaling roles in plants in the phosphorylated form (trehalose-6-phosphate) where it regulates sucrose metabolism and flowering (1, 14-16) and in yeast where it regulates gluconeogenesis and glycolysis (17, 18). Trehalose was further shown to be an autophagy inducer, both in plants and in mammals, with potential biotechnological and clinical implications (19, 20).

In mycobacteria, trehalose is also an essential component of mycolic acids and other cell wall glycolipids, which are major protagonists in *Mycobacterium tuberculosis* pathogenesis (1), the causative agent of the widespread infectious disease tuberculosis. In these organisms, trehalose was further identified as a key signaling molecule of cell-envelope stress playing a role as an activator of the iniBAC operon, which is induced when mycobacteria are exposed to the first line drug isoniazid (21).

All mycobacterial species, with a few exceptions, possess two pathways to synthesize trehalose, the OtsAB and the TreYZ pathway (1). The OtsAB pathway is the most widely distributed trehalose biosynthesis pathway, present in bacteria, archaea and eukaryotes (22, 23). In this pathway, which is conserved and essential in mycobacteria, trehalose is synthesized in a two-step process involving OtsA and OtsB2 enzymes. *M. tuberculosis* mutants showed that the OtsAB pathway was not only the dominant route for trehalose biosynthesis in this pathogen but also required for growth both *in vitro* and in a mouse infection model, with marked growth and virulence defects of OtsA mutants (24) and strict essentiality of OtsB2 due to the toxic effect of trehalose-6-phosphate (T6P) accumulation (24, 25).

OtsA, a glycosyltransferase that belongs to the GT20 family of the CAZY classification (www.cazy.org), uses the α anomer of glucose-6-phosphate (G6P) as acceptor and NDP-glucose as the donor to synthesize T6P, with net retention of the anomeric configuration of the donor substrate (26, 27). Interestingly, OtsAs from different organisms show different donor preferences as shown by kinetic studies and X-ray crystal structures, but the reasons behind these preferences are poorly understood (27-32). Recently, different roles were identified for OtsA beyond its enzymatic activity. OtsA was reported to act as an osmotic stress sensor and morphogenetic protein that can regulate the switch to myceloid growth in *Arthrobacter sp.* strain A3, a pleomorphic soil dwelling actinobacteria (33).

In this work we have purified, crystallized and solved the structure of *Mycobacterium thermoresistibile* OtsA (*Mtr*OtsA). To gain further insight into mycobacterial OtsA, we have obtained structures with substrates, product and pathway product that offer insight for mechanisms of ADP-glucose preference and feedback inhibition by trehalose. We further performed structure-guided point mutations of key residues of the active site and characterized the mutants showing that three mutations are enough to change the donor substrate preference to UDP-glucose. Importantly with these structures, we have also identified a new allosteric site and novel allosteric regulators of this enzyme that link glycolytic and TCA cycle metabolites to the regulation of trehalose synthesis.

## Results

### Overall structure

The structure of *Mtr*OtsA in apo form was solved by molecular replacement, using *E. coli* OtsA structure (PDB code: 1UQU) as the search model. Data collection and refinement statistics are summarized in (Table S1). *Mtr*OtsA is composed of two Rossmann-fold domains with a deep catalytic site at their interface, in an arrangement typical of GT-B glycosyltransferases as described previously for other organisms (27-29). The apo protein crystallized in the I4_1_22 space group, with one protomer per asymmetric unit, and diffracted to ∼1.8 Å resolution. The N-terminal domain is formed by a core of 7 parallel β-strands flanked on both sides by an antiparallel β-strand and surrounded by 8 α-helixes, one of which is composed of the final C-terminal residues (Fig. 1A). The C-terminal domain contains 6 parallel β-strands associated with 9 α-helixes with the last one undergoing a kink and extending to the N-terminal domain, characteristic of the GT-B fold glycosyltransferases (Fig 1A). Analysis of the B-factor distribution shows that the N-terminal domain has the highest values for atomic temperature factors, suggesting that this domain is more dynamic, which is consistent with the large movements of α-1 during catalytic activity.

**Figure 1:**
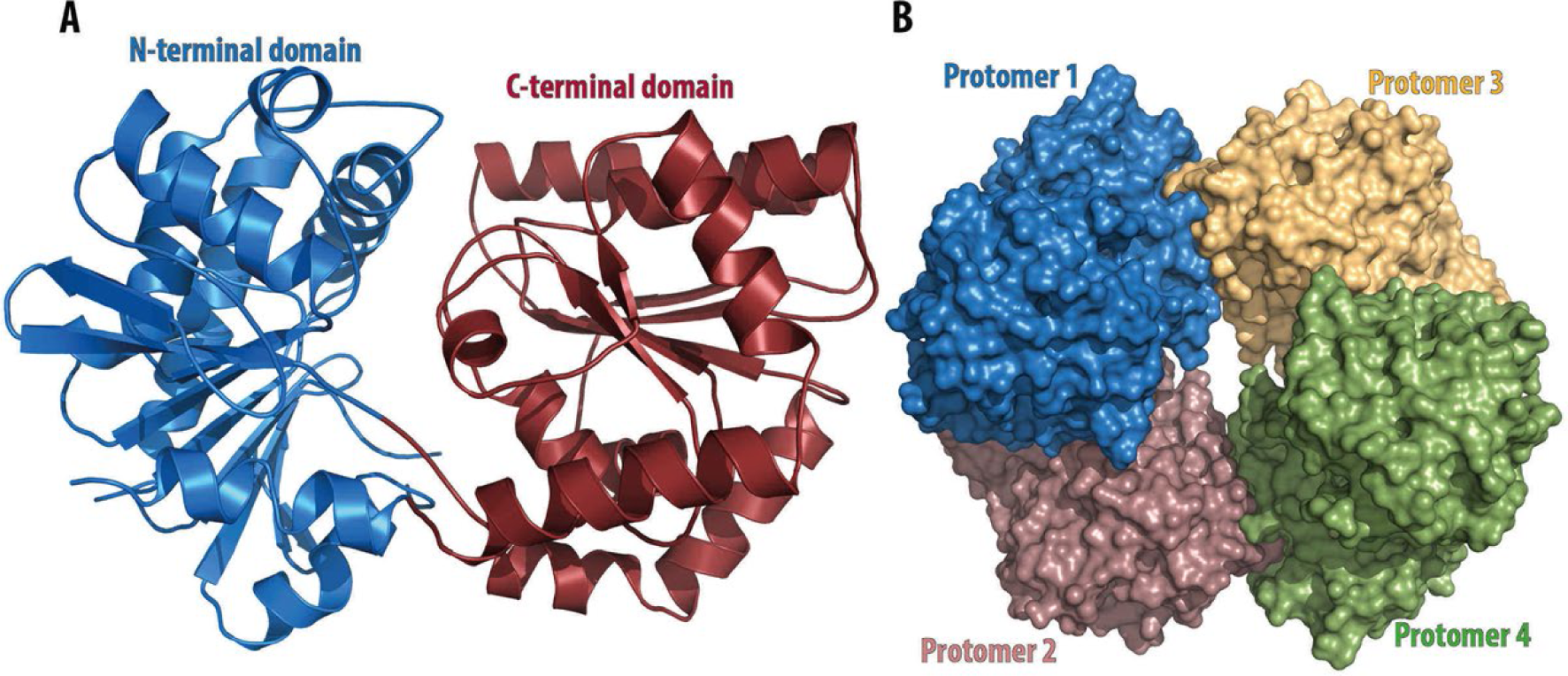
(A) Representation of the overall structure of *M. thermoresistibile* OtsA. The N-terminal domain consists of residues 1-247 and 462-486, the C-terminal domain of residues 248-461. View of *M. thermoresistibile* OtsA tetramer (B).

*Mtr*OtsA forms a tetramer in solution, as previously observed for *M. tuberculosis* OtsA. This tetrameric form is also observed in all the crystal structures reported herein (Fig 1B). The amino acids involved in the tetramer interfaces are not conserved beyond mycobacteria and related species (Fig S1), suggesting that OtsA might have a different molecular assembly in different species. Indeed in *Escherichia coli* both tetrameric and dimeric forms were reported (27) and in *Streptomyces venezuelae, Aspergillus fumigatus* and *Candida albicans* only the dimeric form was observed (28, 29). However, in mycobacteria and closely related organisms the tetramer interfaces are highly conserved, suggesting a tetrameric assembly of OtsA in all of these organisms (Supplementary Fig S1).

### Catalytic site

OtsA, is a glycosyltransferase that uses the α anomer of G6P as acceptor and NDP-glucose as the donor to synthesize T6P. The *Mtr*OtsA catalytic site is located between the two Rossmann-fold domains in a large and deep cavity. Structures with donor substrates were determined by soaking the apo form crystals with ADP-glucose and GDP-glucose (Fig 2). The donor substrates interact primarily with the C-terminal domain through the side chains of the highly conserved Arg286, Lys 291, Asp385, Glu393 and with the side chain of Arg365 (Fig 2), which is conserved in all mycobacteria and closely related species but less conserved outside this group (Fig S1). Backbone interactions with the absolutely conserved residues Gly386, Met387 and Leu389 amine groups are also observed. N-terminal domain interactions with the donor substrate are only observed for His168 when the active site is in an open conformation (Fig 2). To obtain a structure with the acceptor substrate, we co-crystallized OtsA in the presence of 5 mM ADP and G6P. As G6P binds to the protein in the presence of a donor substrate or its nucleotide, the active site adopts to a closed conformation (Fig 3A). The OtsA:ADP:G6P ternary complex crystallized in the P6_2_22 space group, with one protomer per asymmetric unit, and the crystals diffracted up to ∼1.7 Å resolution. In this closed conformation new interactions with the donor substrate are formed with the side chain of Thr42 and the Gly39 peptide-NH function (Fig 3). Due to the absolute conserved nature of Gly39 and the contact it forms with the glucose-bound phosphate, this residue is highly likely to be mechanistically involved in the catalytic activity (27).

**Figure 2:**
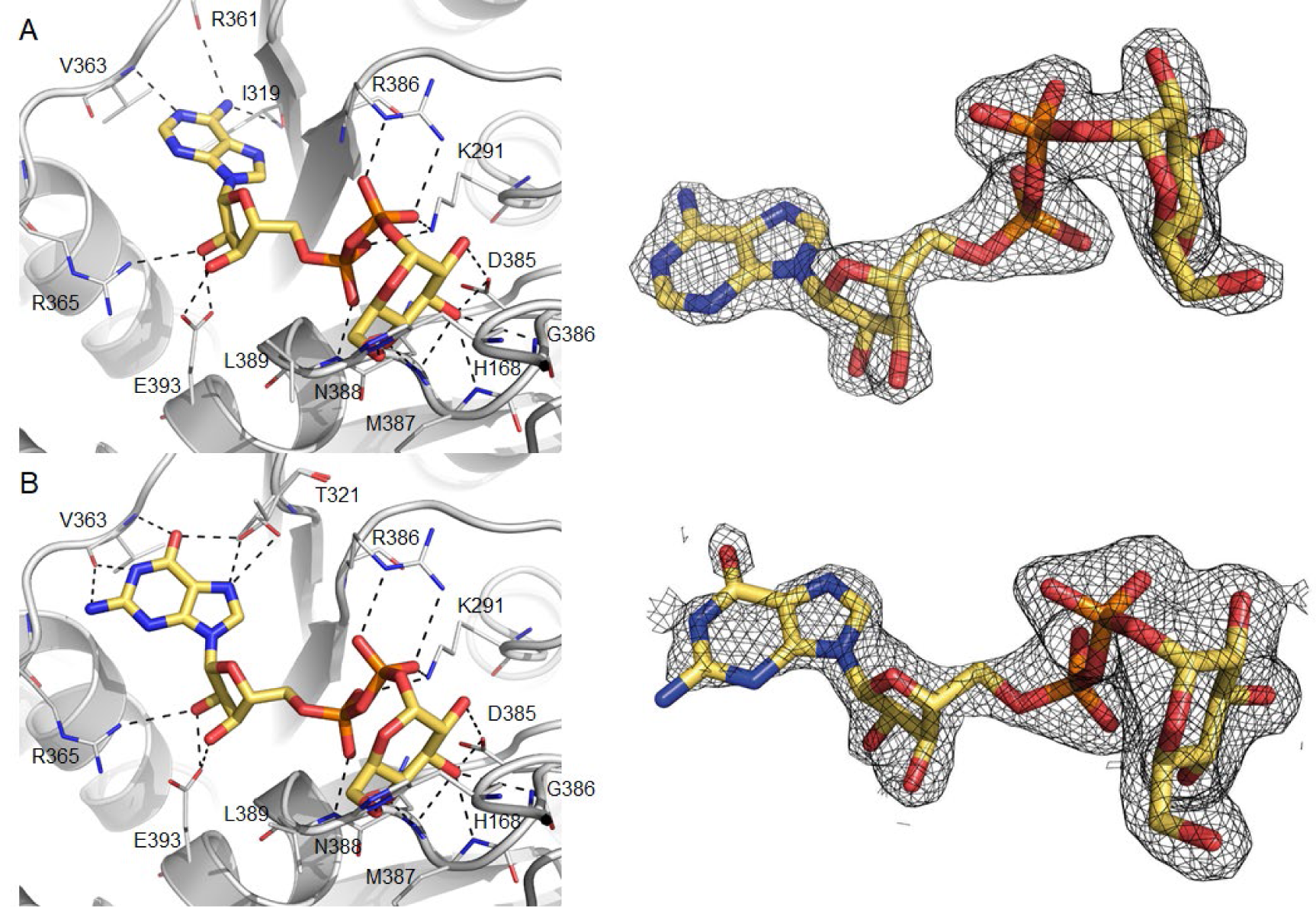
Detailed view of the active site of *M. thermoresistibile* OtsA with ADP-glucose (A) and with GDP-glucose bound (B) with “omit maps” shown. Black dashed lines represent hydrogen bonds.

**Figure 3:**
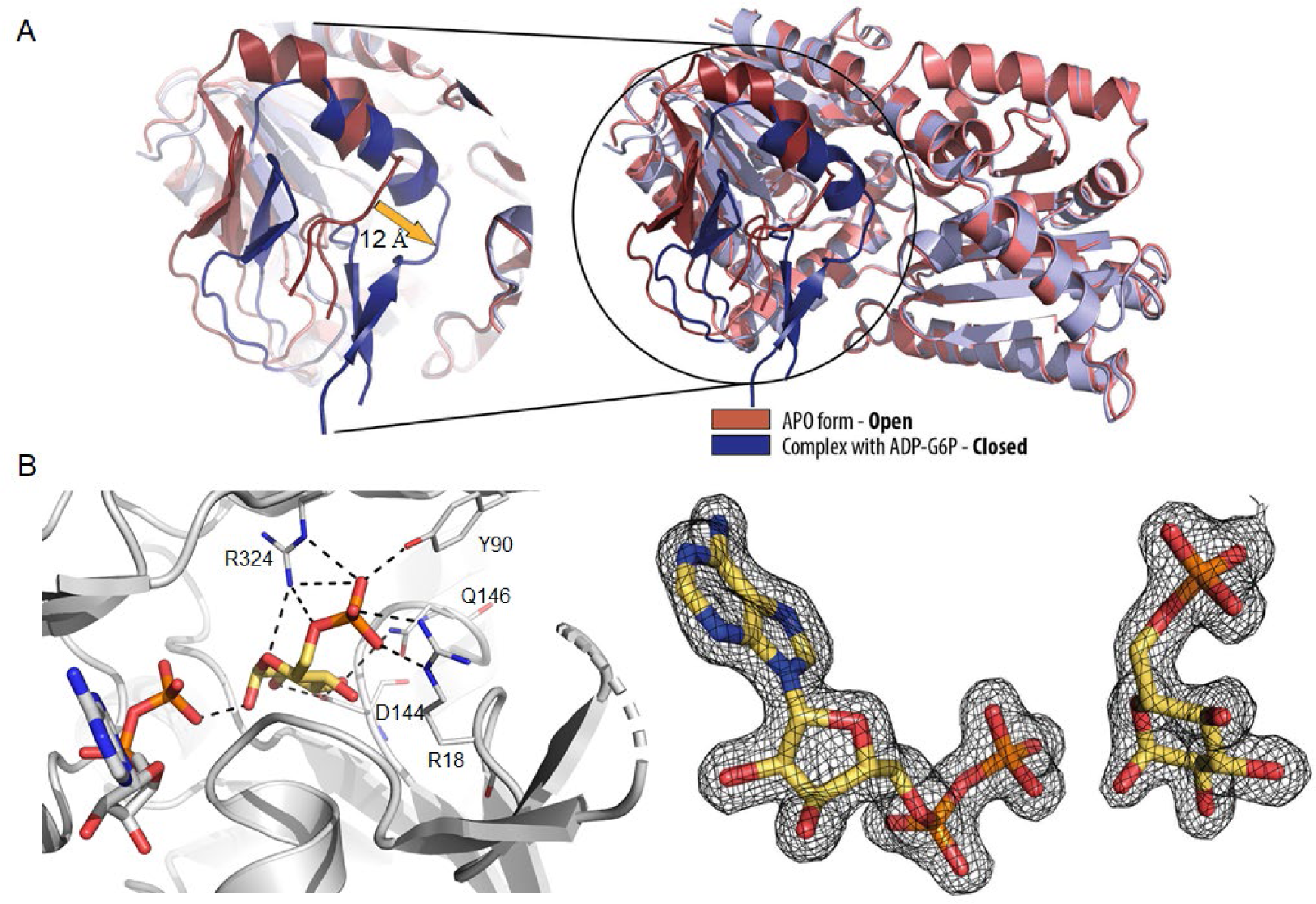
View of OtsA in an open (salmon and red) and closed conformation (violet and blue) and superposition of the two conformations. (B)Representation of the active site of *M. thermoresistibile* OtsA with ADP (white) and G6P (yellow) bound with “omit map” shown. Black dashed lines represent hydrogen bonds.

The acceptor substrate interacts with the highly conserved residues Arg18, Tyr90, Asp144 and Gln146 of the N-terminal domain and with only a single residue of the C-terminal domain, Arg324 (Fig 3B). The residues involved in acceptor substrate interaction are highly conserved in all species (Fig S1). As described for *E. coli* OtsA (27), the OtsA:ADP:G6P ternary structure shows the catalytic site in a closed conformation that substantially differs from the apo form, with α-helix 1 and the Arg35-Gly39 loop region moving up to ∼12 Å (Fig 3A).

### Properties of *M. thermoresistibile* OtsA

*Mtr*OtsA uses ADP-glucose, UDP-glucose and GDP-glucose as glucose donors with decreasing efficiency (Table 1) and G6P as the only acceptor, in accordance to what was reported before for *M. tuberculosis* OtsA (32, 34). The kinetic parameters for *Mtr*OtsA were obtained and are reported in (Table 1) and (Fig. S2) with *K*_*m*_ values for preferred donor substrate (ADP-Glucose) of 0.25 ± 0.02 mM and for the acceptor (G6P) of 3.3 ± 0.1 mM. The *K*_*m*_ for GDP-glucose of 0.29 ± 0.02 mM was in the same range as the one obtained for ADP-glucose, however the turnover was 5 fold lower (Table 1). For UDP-glucose, the enzyme showed a ∼7 fold higher *K*_*m*_ of 1.7 ± 0.1 mM than the one obtained for ADP-glucose (Table 1).

**Table 1:** Kinetic parameters of *M. thermoresistibile* OtsA.

| | Substrate | $K_m$<br>(mM) | $k_{cat}$<br>(s<sup>-1</sup>) | $k_{cat}/K_m$ |
| --- | --- | --- | --- | --- |
| WT | ADP-Glucose | 0.25 ± 0.02 | 26 ± 1 | 104 ± 13 |
|  | GDP-Glucose | 0.29 ± 0.02 | 5.1 ± 0.2 | 18 ± 2 |
|  | UDP-Glucose | 1.7 ± 0.1 | 36 ± 1 | 21 ± 2 |
|  | Glucose-6P | 3.3 ± 0.1 | 26 ± 1 | 7.9 ± 0.5 |
| L319I | ADP-Glucose | 0.71 ± 0.18 | 8.4 ± 0.7 | 12 ± 2 |
|  | GDP-Glucose | 0.35 ± 0.04 | 3.1 ± 0.1 | 8.9 ± 1.4 |
|  | UDP-Glucose | 1.4 ± 0.1 | 8.9 ± 0.4 | 6.4 ± 0.7 |
|  | Glucose-6P | 3.3 ± 0.2 | 6.4 ± 0.2 | 1.9 ± 0.2 |
| V363F | ADP-Glucose | 0.60 ± 0.03 | 11 ± 1 | 18 ± 2 |
|  | GDP-Glucose | 0.74 ± 0.08 | 14 ± 1 | 19 ± 3 |
|  | UDP-Glucose | 0.68 ± 0.06 | 13 ± 1 | 19 ± 3 |
|  | Glucose-6P | 3.1 ± 0.2 | 19 ± 1 | 6.1 ± 0.7 |
| L319I,<br>E367L | ADP-Glucose | 0.63 ± 0.04 | 14 ± 1 | 22 ± 3 |
|  | GDP-Glucose | 0.70 ± 0.05 | 9.2 ± 0.4 | 13 ± 2 |
|  | UDP-Glucose | 0.63 ± 0.03 | 15 ± 1 | 24 ± 3 |
|  | Glucose-6P | 3.7 ± 0.2 | 21 ± 1 | 5.7 ± 0.5 |
| L319I,<br>V363F,<br>E367L | ADP-Glucose | 1.3 ± 0.1 | 19 ± 1 | 15 ± 2 |
|  | GDP-Glucose | 0.90 ± 0.06 | 16 ± 1 | 18 ± 2 |
|  | UDP-Glucose | 0.52 ± 0.05 | 22 ± 1 | 42 ± 6 |
|  | Glucose-6P | 3.2 ± 0.2 | 17 ± 1 | 5.3 ± 0.6 |
| R213E | ADP-Glucose | 0.31 ± 0.01 | 56 ± 2 | 180 ± 13 |
|  | Glucose-6P | 3.3 ± 0.3 | 33 ± 2 | 10 ± 1 |
| R384E | ADP-Glucose | 1.9 ± 0.2 | 3.2 ± 0.1 | 1.7 ± 0.2 |
|  | Glucose-6P | 2.9 ± 0.2 | 2.1 ± 0.2 | 0.77 ± 0.05 |

The preference of ADP-glucose over the other glucose donors was further confirmed by isothermal titration calorimetry (ITC). The binding affinity of substrate donors could only be determined for ADP-glucose in the tested conditions with an observable *K*_*d*_ of 27.17 ± 2.66 µM (Fig S3). Other glucose donors analysed (GDP-glucose and UDP-glucose) had no observable heat of binding and the same was observed for G6P. The lack of heat of binding for GDP-and UDP-glucose is most likely due to their reduced affinity. For G6P case, it reflects a necessity of previous binding of the donor substrate.

### Donor substrate preference is largely mediated by three residues

OtsAs from different organisms have been shown to have different donor substrate preferences, while being capable of using several nucleotide donors (29, 32, 35, 36), which is reflected in lower conservation of the donor substrate interacting residues (Fig S1). Preference for ADP-glucose as the donor substrate in *Mtr*OtsA is conferred by interactions with the deeply buried adenine moiety (Fig 4A). The carbonyl groups of Leu319 and Arg361 interact with the primary amine, and the amide group of Val363 interacts with N1 of the adenine moiety (Fig 4A). A highly coordinated water is also interacting with the primary amine of the adenine moiety and the carbonyl groups of Ala320, Leu359 and Arg361. The guanine moiety of GDP-glucose cannot occupy the same deeply buried pocket as the adenine because its primary amine group would sterically clash with Val363 carbonyl group (Fig 4B). It thus binds to OtsA more weakly than ADP-glucose explaining the substrate preference and consequently the lack of observable heat of binding in the tested ITC conditions. A binary complex structure with OtsA and UDP-glucose was also obtained but the electron density was only observed for glucose and the two phosphates (not shown) indicating a reduced preference for this substrate as confirmed by enzymatic and biophysical data. Comparing mycobacterial OtsAs with *E. coli*, we hypothesised that the ADP preference was mediated by the substitution of an isoleucine (Ile295 in *E. coli*) for a leucine (Leu319 in *M. thermoresistibile*) that allows the primary amine of the adenine moiety to occupy a buried position interacting with the carbonyl group of Leu319 (Fig 4C) that no other nucleotide activated donor can occupy. However, given the residue differences between the two enzymes it is likely that other residues could also play a role. To improve the selection of residues to mutate we employed a computational approach by using a mCSM-lig (37), a software developed by our group that predicts the effects of mutations on binding affinity. The software predicted several that residues within 4.5 Å of the adenine moiety have a destabilizing effect on the interaction with the ligand if mutated to the *E. coli* OtsA equivalent (Table S2). Combining this information with structural analysis we selected three mutations predicted to be destabilizing; Val363Phe, Leu319Ile and Glu367Leu, the latter being a long distance mutation that we hypothesised would have a strong effect on ligand binding.

**Figure 4:**
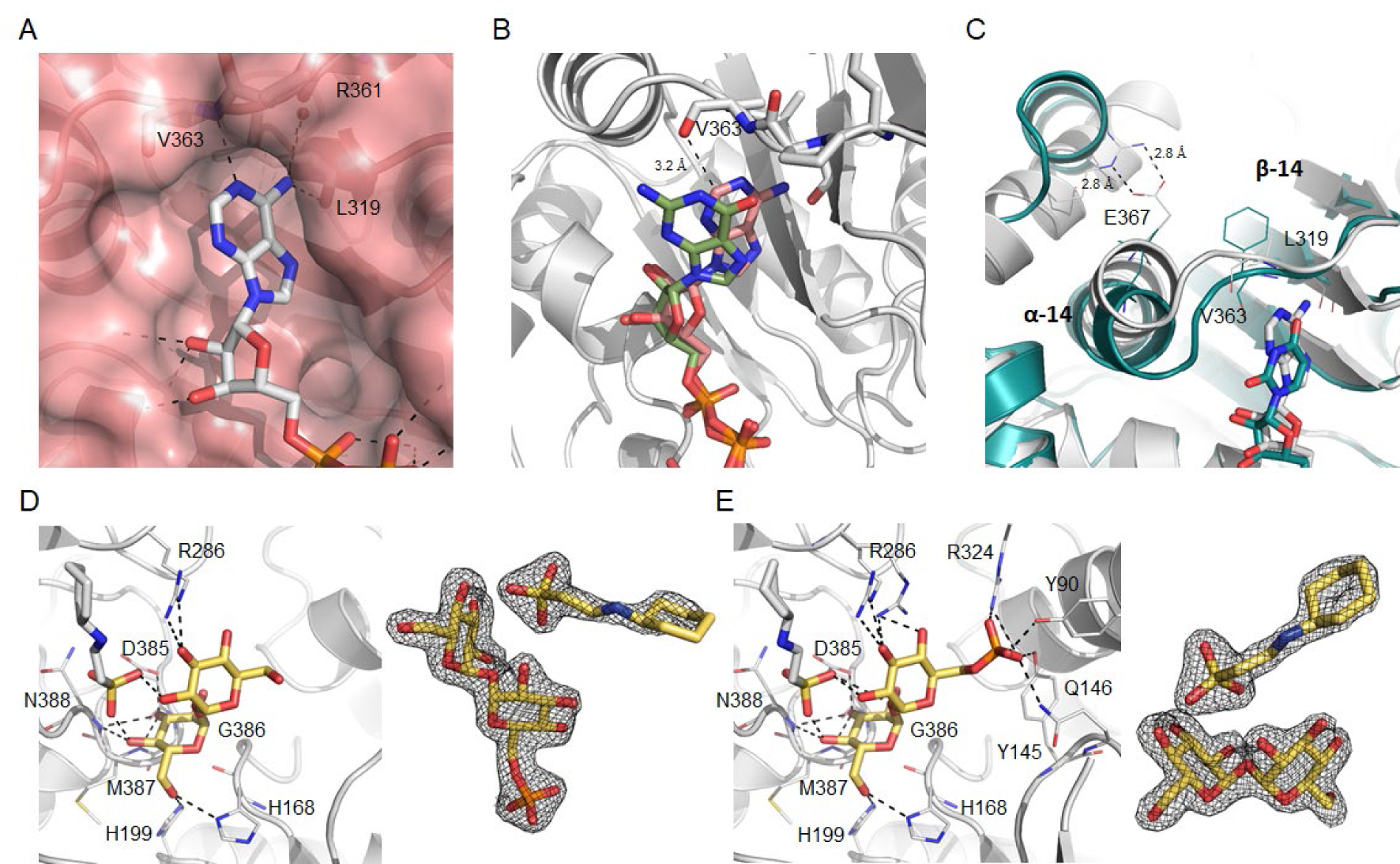
View of the binding site of the adenine moiety of ADP-glucose (A). Superposition of ADP-glucose and GDP-glucose structures (B). Superposition of *Mtr*OtsA structure with ADP-glucose bound and *E. coli* OtsA structure with UDP-glucose bound (PDB code: 1UQU) (C). Alpha helix 14 (α-14) and beta strand 14 (β-14) are shown. Detailed view of the active site of *M. thermoresistibile* OtsA with trehalose (D) and with T6P bound (E) with CHES visible in both structures. “Omit maps” are shown both trehalose, T6P and CHES. Black dashed lines represent hydrogen bonds.

To confirm our hypothesis, we generated several *Mtr*OtsA mutants (Leu319Ile, Val363Phe, Leu319Ile-Glu367Leu and a triple mutant) and we obtained kinetic properties for all of them (Table 1). As we predicted the mutant Leu319Ile showed a ∼3 fold increase e in the *K*_*m*_ for ADP-glucose whereas, *K*_*m*_ was similar GDP-glucose and UDP-glucose when compared to the wild type. Even though GDP-glucose now showed the best *K*_*m*_ among the three different donors, the catalytic efficiency and turnover rate was still higher for ADP-glucose (Table 1). A complete reversal of the donor substrate preference for UDP-glucose was obtained only with a combination of the three mutations (Table 1). The triple mutant showed a ∼5 fold increase in ADP-glucose *K*_*m*_ and a reduction for the UDP-glucose *K*_*m*_ by more than 3 fold (Table 1). The catalytic efficiency was also completely reversed for ADP- and UDP-glucose between the wild-type and the triple mutant (Table 1). Although Leu319Ile showed the largest contribution for ADP-glucose *K*_*m*_ increase, Val363Phe and Glu367Leu were also determinant for the reversal to UDP-Glucose preference (Table 1). This can be explained by the fact that the phenylalanine substitution at position 363 forces the loop between β-14 and α-14 to move towards the nucleotide binding site (Fig 4C), thus establishing stronger interactions with a pyrimidine nucleotide and clashing with purine ones. This effect is also observed for both ADP- and GDP-glucose, with a ∼2.5 fold increase in *K*_*m*_ when compared to the wild type. The leucine substitution at position 367 further helps in this move since the hydrogen bonds between the side chains of Arg266 and Glu367 are no longer present and α-14 is repelled from Arg266 due to the Leu side chain (Fig 4C).

### Feedback inhibition

We soaked both trehalose and T6P into apo OtsA crystals and solved structures with both ligands (Fig 4D and 4E). In both structures OtsA active site assumes an open conformation and two compounds superpose almost perfectly at active site, recapitulating all of the interactions for the donor glucose and also interacting with His199 and Arg286, the latter a residue that interacts with the phosphates of the donor substrate (Fig 4D and 4E). The structure with T6P further shows the phosphate group of the product occupying a similar position to the phosphate group of G6P, interacting with the side chains of Tyr90, Gln146, Arg324 and Tyr145 a residue that does not interact with G6P (Fig 4E). Interestingly the buffer present in the crystallization condition (CHES) is observed in both trehalose and trehalose-6-phosphate bound structures, occupying the site of the nucleotide donor with the sulphate group binding in a phosphate site and interacting with hydroxyl groups from both glucose units of trehalose (Fig 4D and 4E).

After obtaining these structures, it was expected to observe some degree of feedback inhibition of OtsA by both trehalose and T6P. Both trehalose-6-phosphate and trehalose inhibited the enzyme, but the effect of trehalose was more pronounced with ∼40% inhibition at 10 mM while for trehalose-6-phosphate a maximum inhibition of ∼25% was reached at the same concentration (Fig S4). These results show that under physiological conditions OtsA is regulated by trehalose, the final product of the pathway which is highly abundant inside mycobacterial cells, but not by T6P which could not be detected in *M. tuberculosis* cell extracts under physiological conditions (25).

### Identification of an allosteric site in *M. thermoresistible* OtsA

Trehalose is a multipurpose molecule that is abundant in mycobacteria. It can function not only as an energy reserve and as structural component of cell wall glycolipids, but can also lead to the synthesis of glycogen through the TreS-Mak-GlgE pathway and be directly synthesised from glycogen from the TreX-TreY-TreZ pathway. It is therefore likely that trehalose synthesis through the OtsA-B pathway is under strong regulation.

In all the crystallization conditions that contained CHES, we could observe this compound in the crystal structures occupying a pocket formed by the contact interface of 2 OtsA protomers in the tetrameric assembly (Fig 5). The CHES sulphate group directly interacts with the side chains of Arg384 of protomer A and Arg213 of protomer B, forming hydrogen bonds with both, but also hydrophobic hydrogen-π interactions with Phe410 of protomer A (Fig 5). The residues composing this site are completely conserved in mycobacterial and closely related species that are likely to arbour OtsA tetramers but not on others species known to have other oligomeric forms (Fig S1).

**Figure 5:**
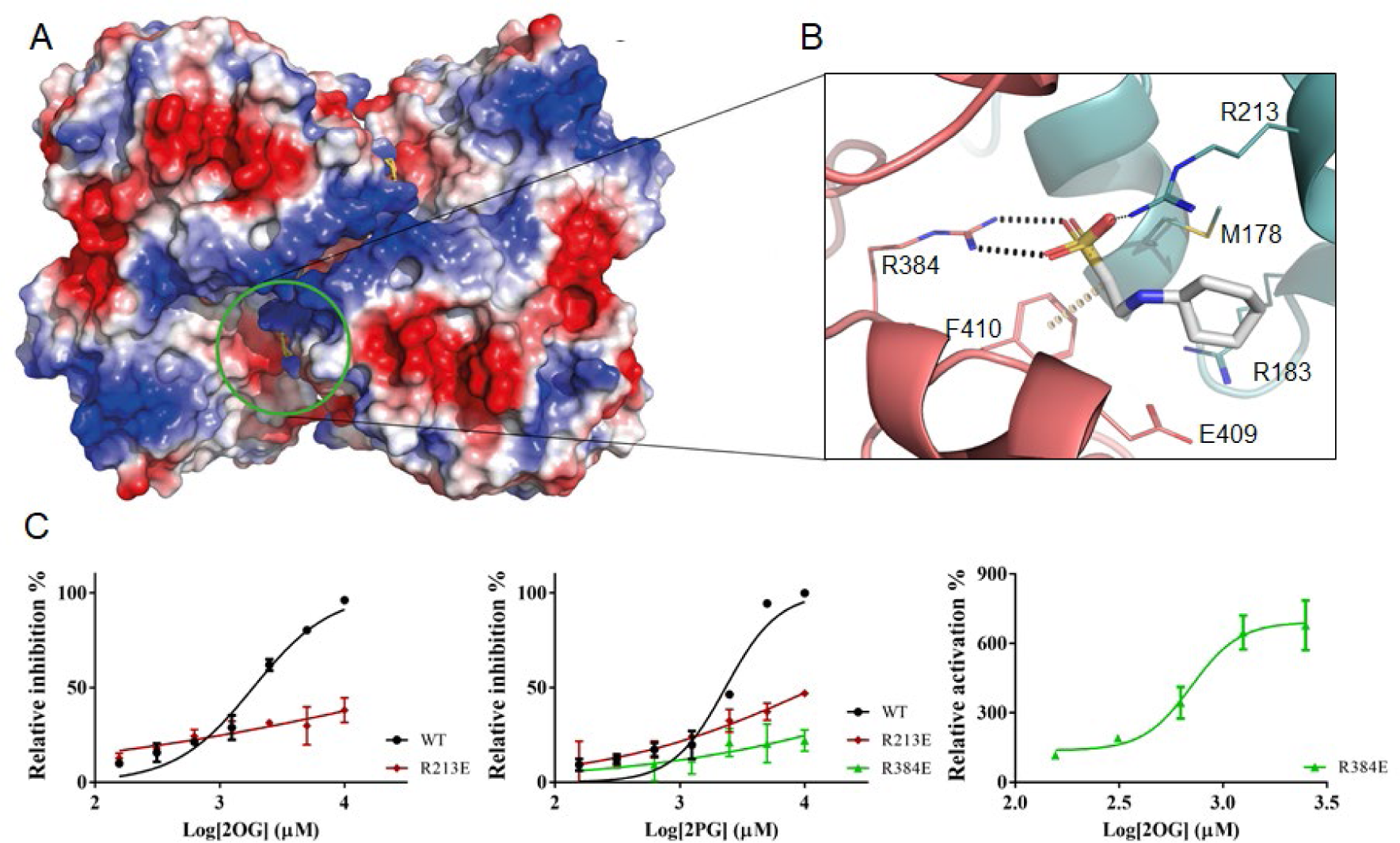
View of the allosteric site of *M. thermoresistibile* OtsA with CHES bound with protein surface electrostatic potential shown (A). Detailed view of the CHES binding site and its interactions with OtsA (B). The interactions were calculated with Arpeggio (60) using the apo structure (5JIJ). Black dots represent hydrogen bonds and yellow disks a carbon-π interaction. The two protomers are colored differently. (C) Activity profiles of *M. thermoresistibile* wild type OtsA and the allosteric site mutants Arg213Glu and Arg384Glu in the presence of the allosteric effectors 2OG and 2PG.

Arg384 sits in a loop that forms extensive contacts with the donor substrate through Asp385, Gly386, Met387 and Leu389 (Fig 2). Ligands interacting with the side chain of Arg384 could therefore have an impact on the activity of the enzyme making this site a prime candidate for allosteric regulation.

While CHES had no discernible impact on OtsA activity in the tested concentrations, it was reported before that F6P acted as an allosteric regulator of *M. tuberculosis* OtsA (34), and the same was reported for yeast OtsA-B complex (38). We have tested *Mtr*OtsA in the same conditions as those reported by Diez and colleagues (34), and could not detect any effect of F6P for *Mtr*OtsA. Although we did not observe any effect of F6P on enzyme activity for *Mtr*OtsAs, we nevertheless attempted to soak this compound in both apo form and OtsA:ADP:G6P ternary complex crystals, and also co-crystallize it in the presence and absence of ADP and G6P, all in CHES free conditions. None of these conditions provided a structure in which F6P occupied an allosteric site. We could however observe F6P when co-crystallized together with ADP, bound to the acceptor substrate site, with OtsA presenting the active site in a closed conformation (Fig S5).

OtsA was found to interact with a multitude of proteins in a large-scale proteomics study (39), including enolase that catalyses the penultimate step of glycolysis. In our search for an allosteric regulator we, therefore, decided to test several glycolytic metabolites glucose-1-phosphate (G1P), fructose-1,6-biphosphate (F16BP), 3-phosphoglycerate (3PG), the enolase substrate 2-phosphoglyceric acid (2PG) and product phosphoenolpyruvate (PEP), but also the master metabolic regulators cAMP and 2-oxoglutarate (2OG) (40), and assess their impact in OtsA activity.

From the several compounds tested only 2OG and 2PG showed clear inhibition of OtsA activity with IC_50_ of 1.8 mM and 2.3 mM respectively (Table 2) and (Fig 5C). To assess whether they could be acting allosterically on *Mtr*OtsA and binding to the site identified by CHES we performed two mutations on the arginines that interacted with CHES (Arg213Glu and Arg384Glu) and tested the two mutants activity in the presence of 2OG and 2PG. The results show that the Arg213Glu mutation abolished the strong response of the protein to 2OG while there was a ∼5 fold increase in 2PG IC_50_ to 12 mM. For the Arg384Glu mutant 2OG becomes unexpectedly becomes a strong activator with an EC_50_ of 0.7 mM while 2PG has no effect (Table 2).

**Table 2:** Effect of allosteric regulators in *M. thermoresistibile* OtsA activity. Values are in (mM) and 95% confidence intervals are given in brackets.

|  | WT | R213E | R384E |
| --- | --- | --- | --- |
| 2OG | IC<sub>50</sub> = 1.8<br>(1.6-2.1) | IC<sub>50</sub> = 73<br>(18-302) | EC<sub>50</sub> = 0.70<br>(0.56-0.88) |
| 2PG | IC<sub>50</sub> = 2.3<br>(2.0-2.7) | IC<sub>50</sub> = 12<br>(6.3-24) | IC<sub>50</sub> < 100 |

## Discussion

Trehalose, an essential disaccharide in mycobacteria, is one of the critical components of the cell wall. The OtsA-B pathway is an essential source of trehalose for these organisms with knockouts of this pathway being growth defective or non-viable (24, 25). Here we reveal the first mycobacterial OtsA structure. The overall structure of *M. thermoresistibile* OtsA is similar to the reported structures of *E. coli, S. venezuelae* and *Paraburkholderia xenovorans* OtsAs despite relatively low sequence identity (30-35%). Remarkably, OtsAs have also a highly similar fold to the pseudo-glycosyltransferases VldE and ValL involved in validamycin A synthesis, a potent antifungal agent (41, 42).

Nevertheless, there are substantial differences in the oligomeric organization of bacterial OtsAs, with tetrameric, dimeric and protomeric forms reported. While mycobacteria and related species present conservation of interfaces showing that the observed tetrameric form for *Mtr*OtsA is present in all these organisms, beyond this group such conservation is absent, which is consistent with other oligomeric states (Figure S1).

The structures obtained in this work allow us to explain how mycobacterial OtsA preference for ADP-glucose, is conferred by Leu319 which creates a cavity to accommodate the primary amine of the adenine moiety allowing it to interact with Leu319 and Arg361carbonyl groups. It is further mediated by Val363 carbonyl group that does not allow GDP-glucose to occupy the same position and Glu367. Mutating these three residues to the *E. coli* OtsA equivalents completely changed the preference to UDP-glucose, the preferred *E. coli* OtsA donor substrate. Both Leu319 and Glu367 are highly conserved in all mycobacterial and closely related species analysed but not beyond this group, while Val363 can be replaced by other small hydrophobic amino acids such as alanine and isoleucine in other mycobacterial species and closely related species (Figure S1). However, outside this group Val363 is replaced by bulkier residues such as tyrosine and phenylalanine (Figure S1). These differences are suggestive of alternative donor substrate preferences for organisms outside the sub-order *Corynebacterineae*.

Trehalose is not only abundant inside the mycobacterial cell but also involved in a cycle that links it to the synthesis and degradation of glycogen (43). We have shown that OtsA is feedback inhibited by trehalose but not by T6P even though the structures obtained show that both compounds form extensive interactions with OtsA. However, given that these two structures were obtained by soaking apo form crystals with high concentrations of trehalose and T6P, and that CHES is also observed at the active site interacting with both compounds, it is possible that these two structures do not entirely reflect their natural interactions with the enzyme. The toxic accumulation of T6P in OtsB2 knockout mutants (25) can be explained by these results since OtsA shows very low sensitivity to T6P. Furthermore trehalose levels are reduced in the mutant (25) further contributing to an increase in OtsA activity leading to higher T6P production.

Regulation of OtsA by phosphorylation and methylation has been observed for yeast trehalose-6-phosphate synthase complex (44, 45) and phosphorylation of *E. coli* OtsA has also been reported (46) in two residues close to the active site. Even though only one of the residues is conserved in mycobacteria (Ser323 for *Mtr*OtsA), phosphorylation of this residue would most likely lead to inactivation of the enzyme due to the location close to the acceptor binding site (Fig S6). Interestingly this residue is highly conserved in all sequences analysed with a single exception where it is substituted by a threonine (Figure S1), suggesting that phosphorylation of this residue might be a common regulatory mechanism of OtsA activity.

Allosteric regulation was reported for both *M. tuberculosis* OtsA and yeast trehalose phosphate synthase complex, mediated by F6P (34, 38). The structure obtained with F6P shows this compound occupying the donor site, in the presence of ADP, with the enzyme adopting a closed conformation. Nevertheless, OtsA could only use G6P as an acceptor and F6P showed no effect in the enzyme activity, indicating that the presence of F6P in the acceptor site was forced by the high concentration used for co-crystallization (5mM). The observed differences in regulation by F6P between mycobacterial enzymes are difficult to reconcile since the allosteric site is highly conserved in mycobacterial OtsAs and closely related species (Fig S1).

*Mtr*OtsA is allosteric regulated by 2PG, the substrate of enolase, a glycolytic enzyme, and 2OG a master metabolic regulator that sits at the interface of carbon and nitrogen metabolism. The observed effect is within relevant physiological concentrations for both compounds (47, 48).

The effect of 2PG on OtsA and the reported interaction with enolase suggests an interplay between these two enzymes that would be interesting to explore further on the enolase side. Enolase is a multifunctional protein involved in a variety of cellular processes, beyond its enzymatic activity, that include response to oxidative and thermal stress (49), and is further involved in the RNA degradosome activity, by regulating RNA stability in response to stress (50, 51). Given that trehalose is a known chemical chaperone and compatible solute (1), the association between enolase and OtsA, through the effect of 2PG in OtsA activity and the reported interaction between the two enzymes, points to further regulation to stress response that would be interesting to explore further on the enolase side.

2OG is a molecule that sits at the interface of carbon and nitrogen metabolism and that has been shown to regulate many different pathways (40). The role of trehalose as an energy reserve molecule (1) and its relationship with synthesis and degradation of glycogen (52) points to a mechanism in which 2OG influences the synthesis of these two compounds through regulation of OtsA activity that deserves to be further explored.

We have shown that OtsA can be allosteric inhibited, however a single mutation at the allosteric site changed the behaviour of 2OG, but not 2PG, from an inhibitor to an activator. This hints at the possible existence of allosteric activators of OtsA to be discovered.

The conservation of allosteric site residues and oligomeric assembly in the sub-order *Corynebacterineae* but not outside, suggests that allosteric regulation of OtsA through this site might be limited to this group of organisms. The results of this work are a significant step forward in understanding the regulation of trehalose synthesis in mycobacteria the structural reasons behind substrate preference, and provide new important insight into this enzyme.

## Methods

### Bacterial strains and cloning

*M. thermoresistibile* (DSM 44167) *otsA* gene was amplified from chromosomal DNA obtained from the Deutsch Sammlung von Mikroorganismen und Zellkulturen GmbH (DSMZ, Braunschweig, Germany). Primers were designed based on the sequence available in NCBI database and the gene was cloned between the BamHI and HindIII sites in pET28a vector (Novagen), modified with an N-terminal 6xHis-SUMO tag. The resulting plasmid was confirmed by DNA sequencing and transformed into *E. coli* BL21(DE3) strain (Invitrogen). Six *Mtr*OtsA mutants (Arg213Glu, Leu319Ile, Val363Phe, Arg384Glu, Leu319Ile-Glu367Leu and Leu319Ile-Val363Phe-Glu367Leu) were also constructed by site-specific mutagenesis, sequenced and transformed into *E. coli* BL21(DE3) strain. Primers used in this work are listed in table S3.

### Recombinant expression and protein purification

Transformed *E. coli* BL21(DE3) cells were grown to mid-exponential growth phase (OD_610_ = 0.6) in LB media (Invitrogen) containing 30 mg L^-1^ kanamycin at 37 °C. Isopropyl β–D-1-thiogalactopyranoside (IPTG) was then added at a final concentration of 0.5 mM to induce gene expression and the temperature was lowered to 18 °C. Cells were harvested 18 h-20 h later by centrifugation and re-suspended in 20 mM TRIS pH 7.5, 500 mM NaCl and 20 mM Imidazole with protease inhibitor tablets (Roche), DNAseI and 5 mM MgCl_2_. Cells were lysed by sonication and cell lysate was centrifuged at 27000 g for 30 min to remove cell debris.

Recombinant *Mtr*OtsA was purified with a HiTrap IMAC Sepharose FF column (GE-Healthcare), equilibrated with 20 mM TRIS pH 7.5, 500 mM NaCl and 20 mM Imidazole. Elution was performed in the same buffer, but with 500 mM Imidazole. Imidazole was removed with a desalting column and SUMO tag was cleaved overnight at 4 °C by adding Ulp1 protease at 1:100 ratio in 20 mM TRIS pH 7.5, 500 mM NaCl. SUMO tag, Ulp1 protease and uncleaved SUMO-OtsA were removed with a HiTrap IMAC Sepharose FF column (GE-Healthcare), equilibrated with 20 mM TRIS pH 7.5, 500 mM NaCl and 20 mM imidazole. Flow through containing OtsA was collected, concentrated and loaded to a Superdex 200 column equilibrated with 20 mM TRIS pH 7.5, 500 mM NaCl. Fraction purity was determined by SDS-page and purest fractions were pooled concentrated to ∼10 mg.ml^-1^ in 20 mM TRIS pH 7.5 350 mM NaCl, flash frozen in liquid nitrogen and stored at −80 °C. The same purification protocol was used for all *Mtr*OtsA mutants.

### Crystallization and data collection

*Mtr*OtsA crystallization screens and optimization were performed at 18 °C using sitting-drop vapour diffusion method. 300 nl of pure OtsA at 10 mg.ml^-1^ was mixed in 1:1 ratio with well solution using a Phoenix robot (Art Robbins). Initial conditions were obtained in classics suite crystallization screen (Qiagen), solution 36. Crystals obtained in this condition diffracted only up to 3 Å. Therefore, further optimization was performed using the additive screen HT (Hampton Research) and ethylene glycol was found to be the best additive. The final optimized condition consisted of 0.7 M sodium potassium tartrate, 0.1 M CHES pH 10 and 10% v/v ethylene glycol. Crystals appeared after 4 days in this condition. To obtain ligand-bound structures soaking was performed in the optimized condition using the hanging drop vapour diffusion method as follows: 1 µl of protein storage buffer containing 5 mM of ligand was mixed with 1 µl of reservoir solution and drops were left to equilibrate against 500 µl of reservoir solution for 3 days. Crystals were then transferred to the pre-equilibrated drops and incubated for 24h. A cryogenic solution was prepared by adding ethylene glycol up to 25% v/v to mother liquor. Crystals were briefly transferred to this solution, flash frozen in liquid nitrogen and stored for data collection. To obtain an ADP-G6P-OtsA ternary complex co-crystallization with 5mM ADP and G6P was performed instead, as all attempts to soak G6P alone or G6P in the presence of ADP have failed. Crystals were obtained in the Wizard Classic I&II screen (Rigaku), solution E10 and were flash frozen in liquid nitrogen after a brief soak in a solution containing mother liquor and 25% ethylene glycol. The same condition was used to obtain ADP:F6P:OtsA complex.

All data sets were collected at stations I04, I04-1 and I24 at Diamond Light Source (Oxford, UK). Data collection and refinement statistics are summarized in (Table S1).

### Structure solution and refinement

Diffraction data were processed and reduced using MOSFLM (53) and Aimless (54) from the CCP4 suite (55), or autoPROC from Global Phasing Limited (56). The apo form crystalized in I4_1_22 space group with one protomer per asymmetric unit. ADP-G6P-OtsA ternary complex crystallized in P6_2_22 space group again with one protomer per asymmetric unit.

Initial phases were determined with PHASER (57) from PHENIX software package (58) using the structure of *E. coli* OtsA (PDB entry 1UQU) (31) as a search model. Model building was done with Coot (59) and refinement was performed in PHENIX (58). Structure validation was performed using Coot and PHENIX tools (58, 59). All the figures were prepared with Pymol (http://www.pymol.org).

### Prediction of mutations in ligand affinity

To provide an insight on which residues to mutate, we used mCSM-lig a software that predicts the effect of mutations in ligand affinity (37) on the X-ray crystal structure of OtsA with ADP-glucose (5JIO).

### Enzymatic Assays

Formation of T6P was assessed by a continuous colorimetric assay that followed the release of NDP by measuring the oxidation of NADH at 340nm, in the presence of pyruvate kinase and lactate dehydrogenase. All reagents used were obtained from Sigma-Aldrich. The enzymatic reactions (200 µl) were performed at 37 °C and contained 50 mM TRIS pH 7.5, 200 mM NaCl, 10 mM MgCl_2_, 50 mM KCl, 0.3 mM NADH, 2.5 mM phosphoenolpyruvate (PEP), 10 units of pyruvate kinase/lactate dehydrogenase, 0.1 µM enzyme and varying concentrations of G6P and NDP-glucose to determine kinetic parameters.

The effect of trehalose and T6P as feedback inhibitors, and G1P, F6P, F16BP, CHES, 2PG, 3PG, cAMP and 2OG as allosteric regulators, was tested in the conditions described above but with fixed concentrations of G6P and ADP-glucose (both 0.3 mM). The same conditions were used to test possible allosteric regulators for wild type OtsA and mutants Arg213Glu and Arg384Glu. All experiments were performed in triplicate in a PheraStar plate reader (BMG Labtech) and data was analysed with Prism 5 (Graphpad Software). To assess the inhibitory effect of ADP and allosteric effect of phosphoenolpyruvate an end-point assay was performed instead. Reactions (100 µl) containing 50 mM TRIS pH 7.5, 200 mM NaCl, 10 mM MgCl_2_, 50 mM KCl, 4 µM *Mtr*OtsA, 0.3 mM ADP-glucose, 0.3 mM G6P and varying concentrations of ADP and PEP were incubated at 37 °C and stopped at different time points with 10 µl of 1 M HCl, incubated for 1 minute and neutralized with NaOH. The reactions were diluted with 100 µl acetonitrile and run at 25 °C on a Waters I-class UHPLC with a PDA detector (258 nm) using a ACQUITY UHPLC BEH Amide column (2.1 × 100 mm, 1.7 μm). Gradient elution (delivered at 0.13 mL/min) was employed using 80/20 acrilamide/water with 0.1 % NH_2_OH (A) and water with 0.1 % NH_2_OH (B) which started at 90 % A and decreased linearly to 55 % A over 10 min.

### Isothermal Titration Calorimetry

Binding interaction between OtsA and ligands was characterized at 25 °C, using a Microcal ITC200 titration calorimeter (Microcal). OtsA concentration of 60 µM was used for all titrations. Ligands (1 mM) were injected in 2ul aliquots. Titration data was recorded in 20mM TRIS pH 7.5, 500mM NaCl. Data was analysed by fitting a simple single-site model using Origin software (Microcal).

