## Supplemental data for "Mycobacterial OtsA structures unveil substrate preference mechanism and allosteric regulation by 2-oxoglutarate and 2-phosphoglycerate"

Tom L. Blundell

Address: Department of Biochemistry, University of Cambridge, 80 Tennis Court Road, CB2 1GA Cambridge, UK.

Vitor Mendes

Address: Department of Biochemistry, University of Cambridge, 80 Tennis Court Road, CB2 1GA Cambridge, UK.

### Results

**Table S1:** X-ray crystallography data collection and final refinement statistics

| Ligand# | **APO** | **ADP-glucose** | **GDP-glucose** | **ADP G6P** | **ADP-F6P** | **Trehalose** | **T6P** |
|---|---|---|---|---|---|---|---|
| PDB ID | 5JIJ | 5K41 | 5K42 | 5JIO | 5L3K | 5K5C | 5K44 |
| **Data collection*** | | | | | | | |
| Space group | $I4_122$ | $I4_122$ | $I4_122$ | $P6_222$ | $P6_2$ | $I4_122$ | $I4_122$ |
| Cell parameters: | | | | | | | |
| a [Å] | 126.86 | 127.41 | 127.31 | 105.36 | 216.96 | 127.32 | 127.23 |
| b [Å] | 126.86 | 127.41 | 127.31 | 105.36 | 216.96 | 127.32 | 127.23 |
| c [Å] | 207.19 | 205.62 | 206.78 | 158.87 | 159.84 | 207.38 | 207.61 |
| α/β/γ [˚] | 90/90/90 | 90/90/90 | 90/90/90 | 90/90/120 | 90/90/120 | 90/90/90 | 90/90/90 |
| Resolution range [Å] | 60.66 – 1.82 | 102.81 – 1.97 | 108.41 – 1.92 | 91.24– 1.71 | 159.84 – 2.31 | 103.69 – 1.85 | 103.80 – 1.93 |
| | (1.86 – 1.82) | (2.20 – 1.97) | (2.15 – 1.92) | (1.87 – 1.71) | (2.43 – 2.31) | (2.02 – 1.85) | (2.15 – 1.93) |
| No. of observations | | | | | | | |
| total | 833006 | 753000 | 842990 | 1030194 | 1927301 | 923936 | 817022 |
| | (50109) | (216413) | (231847) | (193800) | (266711) | (184758) | (229455) |
| unique | 75576 | 59763 | 64773 | 56816 | 187701 | 72774 | 64496 |
| | (4427) | (16773) | (18176) | (13323) | (27326) | (17204) | (18114) |
| $R_{merge}$ | 0.116(0.716) | 0.154 (0.669) | 0.078 (0.958) | 0.096 (0.790) | 0.268 (1.138) | 0.076 (0.762) | 0.093 (0.544) |
| I/σ(I) | 9.5 (2.0) | 9.5 (2.8) | 23.1 (3.3) | 19.1 (3.3) | 7.0 (2.1) | 20.1 (3.0) | 15.6 (3.5) |
| Completeness [%] | 100.0 (100.0) | 100.0 (99.9) | 100.0 (99.9) | 100.0 (100.0) | 99.9 (99.9) | 100.0 (100.0) | 100.0 (100.0) |
| Multiplicity | 11.0 (11.3) | 12.6 (12.9) | 13.0 (12.8) | 18.1 (14.5) | 10.3 (9.8) | 12.7 (10.7) | 12.7 (12.7) |
| **Refinement** | | | | | | | |
| Refinement program | PHENIX | PHENIX | PHENIX | PHENIX | PHENIX | PHENIX | PHENIX |
| Resolution [Å] | 60.66 – 1.82 | 90.09 – 1.97 | 45.01 – 1.92 | 91.24 – 1.71 | 121.74 – 2.31 | 41.58 – 1.85 | 40.23 – 1.93 |
| No. reflections | 74139 | 59634 | 64577 | 56756 | 187654 | 72668 | 64406 |
| $R_{work}/R_{free}$ [%] | 16.5/18.6 | 16.5/19.1 | 15.6/18.4 | 15.2/17.9 | 15.3/21.3 | 15.1/17.5 | 14.9/17.4 |
| RMS deviations | | | | | | | |
| Bonds [Å] | 0.007 | 0.007 | 0.007 | 0.007 | 0.009 | 0.007 | 0.008 |
| Angles [˚] | 1.009 | 1.042 | 1.045 | 1.092 | 1.048 | 1.028 | 1.030 |
| Ramachandran | | | | | | | |
| Favoured [%] | 97 | 97 | 97 | 97 | 95 | 97 | 96 |
| Outliers [%] | 0 | 0 | 0 | 0 | 0 | 0 | 0 |

* Parameters shown in brackets are for the highest resolution shell

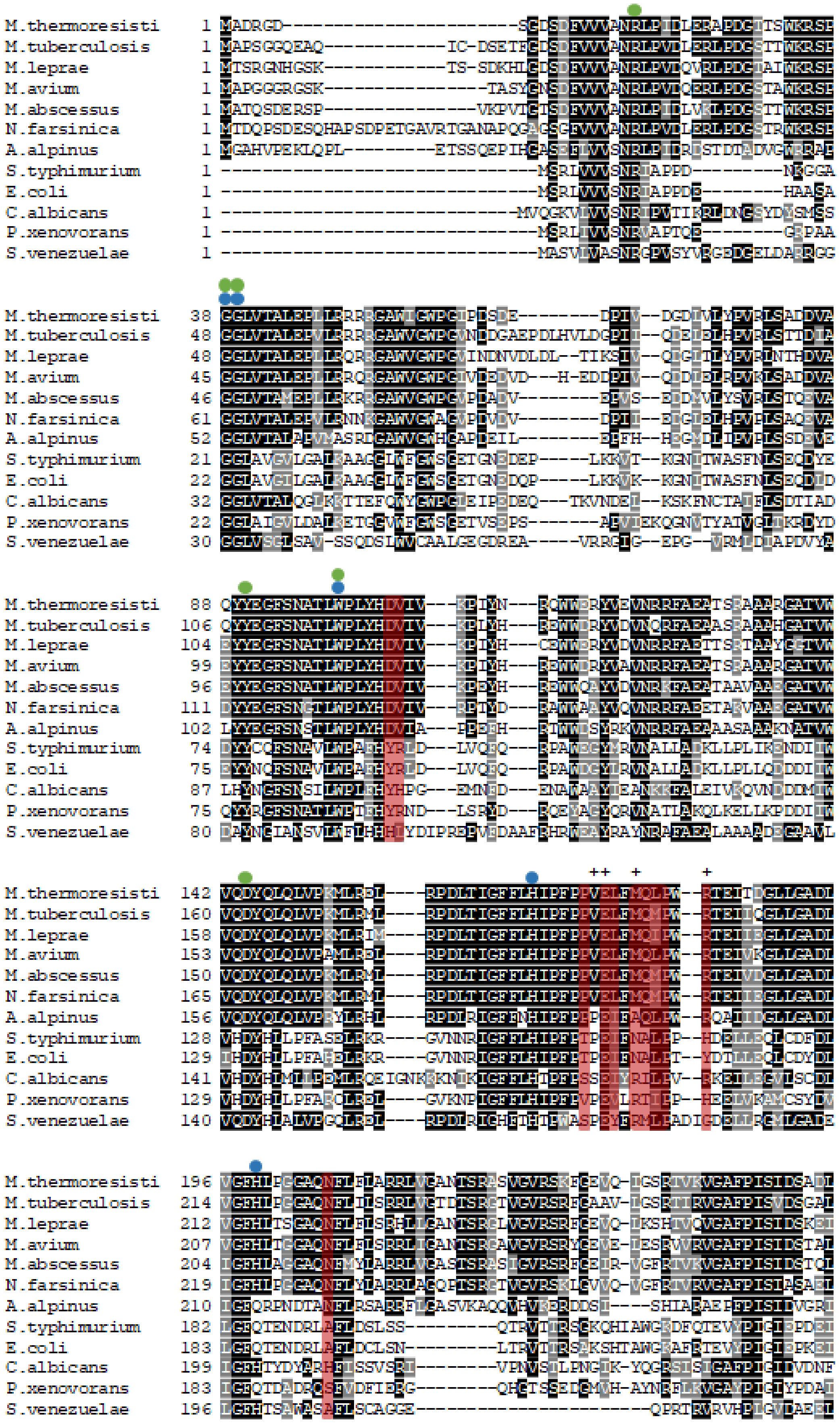
M.thermoresisti
M.tuberculosis
M.leprae
M.avium
M.abscessus
N.farsinica
A.alpinus
S.typhimurium
E.coli
C.albicans
P.xenovorans
S.venezuelae

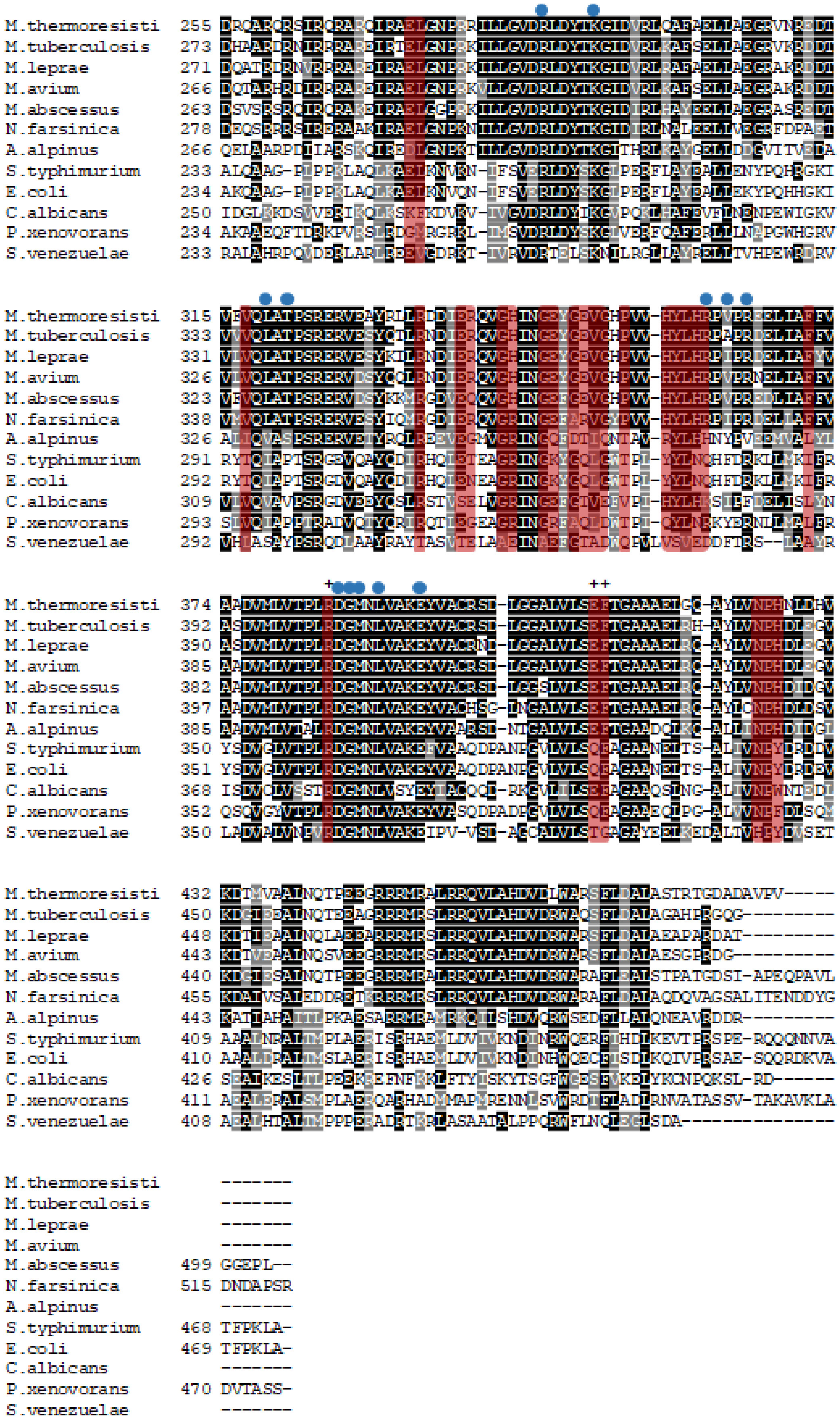
M.thermoresisti
M.tuberculosis
M.leprae
M.avium
M.abscessus
N.farsinica
A.alpinus
S.typhimurium
E.coli
C.albicans
P.xenovorans
S.venezuelae

**Supplementary Figure S1:** Sequence comparison of OtsA from *Mycobacterium thermoresistibile*, *Mycobacterium tuberculosis*, *Mycobacterium leprae*, *Mycobacterium avium*, *Mycobacterium abscessus*, *Nocardia farcinica*, *Arthrobacter alpinus*, *Salmonella typhimurium*, *Escherichia coli*, *Candida albicans*, *Paraburkholderia xenovorans* and *Streptomyces venezuelae*. Residues that contact with the substrates are highlighted with blue circles (donor site) and green circles (acceptor site). The allosteric site residues are marked with crosses and tetramer interfaces are highlight in red. The tetramer interfaces are highly conserved only in the several mycobacteria and *N. farcinica* and less *A. alpinus*. The remaining non-actinobacterial species show very little conservation of the interfaces. The same is observed for the allosteric site. Acceptor site residues are conserved throughout. Donor site residues are less conserved indicating the known differences in substrate preference. The multiple sequence alignment was performed with T-Coffee (1).

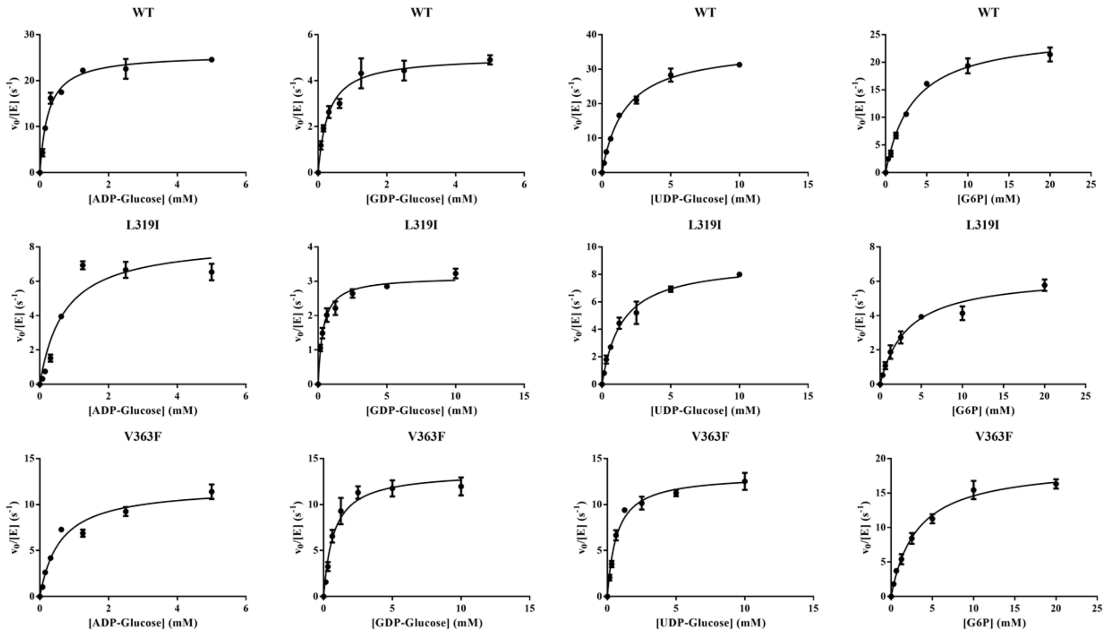

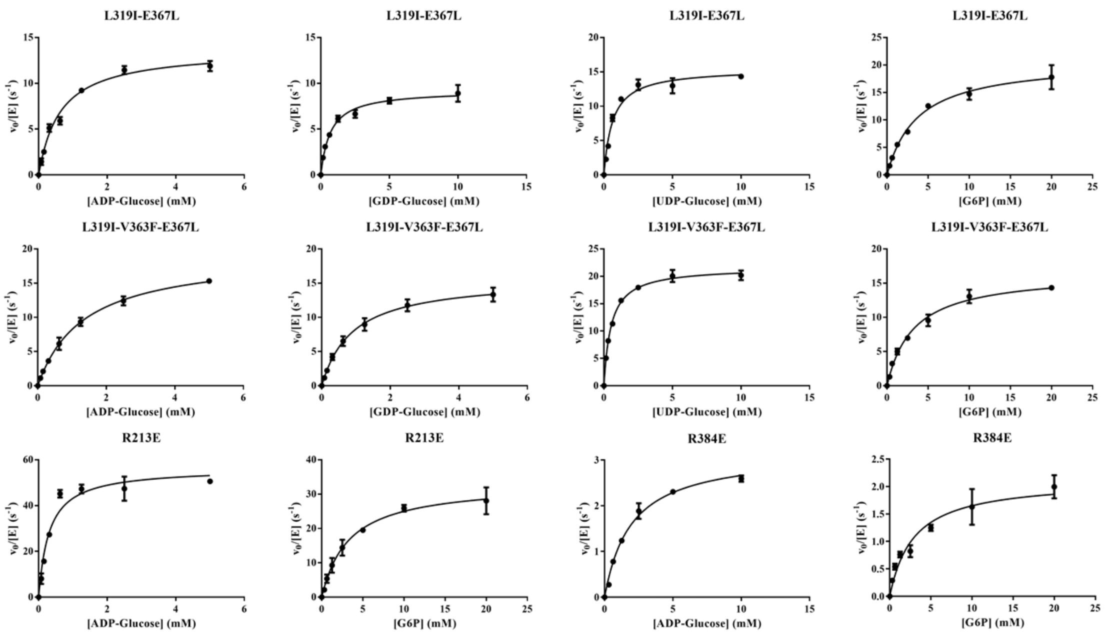


**Supplementary figure S2:** *Mtr*OtsA wt and mutants kinetics. G6P concentration was kept at 10 mM for all NDP-glucose $K_m$. ADP-glucose concentration was kept at 2.5 mM for G6P $K_m$.

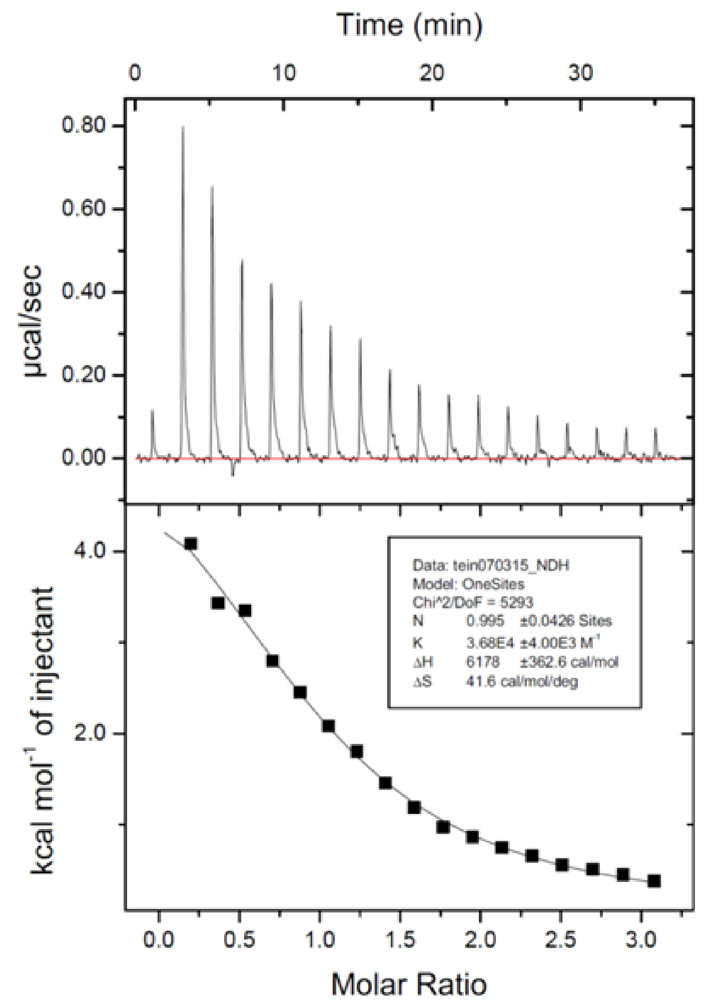


**Supplementary figure S3:** ITC trace with *Mtr*OtsA for ADP-glucose

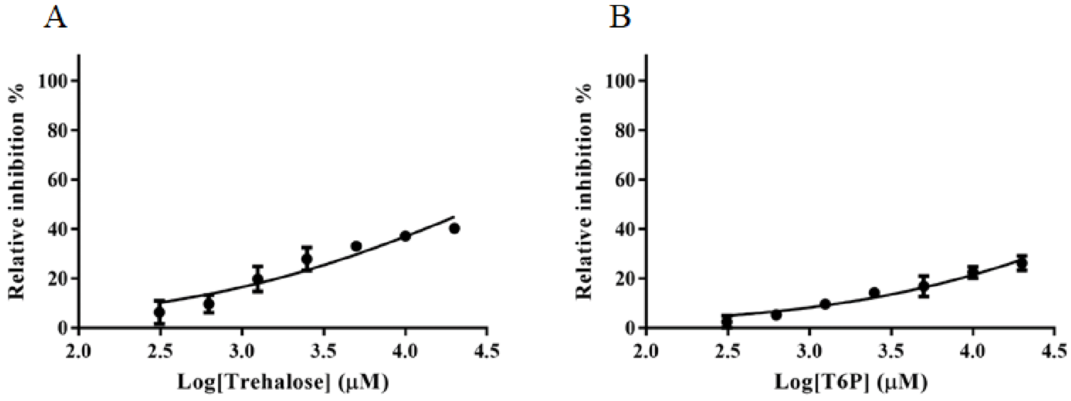


**Supplementary figure S4:** Feedback inhibition by trehalose (A) and T6P (B).

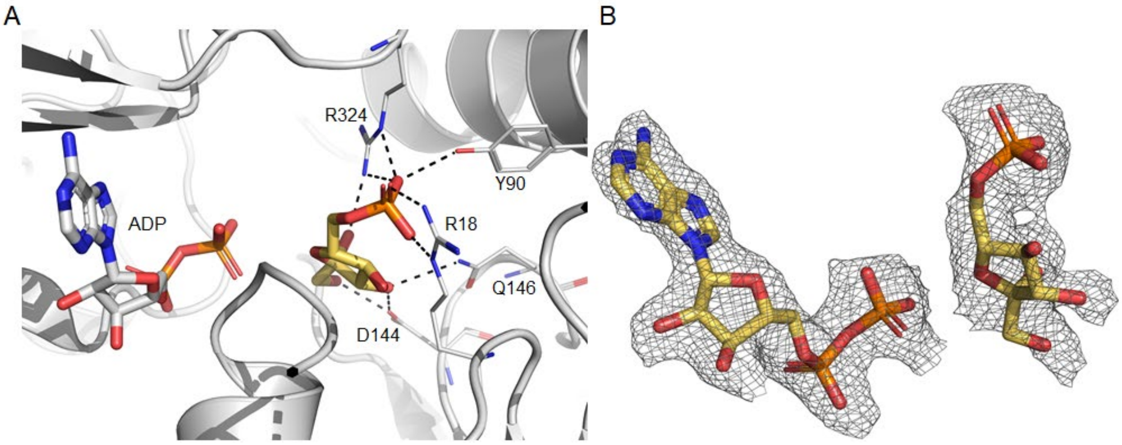


**Supplementary Figure S5:** (A) - View of the active site of *M. thermoresistibile* OtsA complexed with ADP (white) and F6P (yellow), with F6P occupying the acceptor site. “Omit” map for the two ligands is shown in (B). Black dashed lines represent hydrogen bonds.

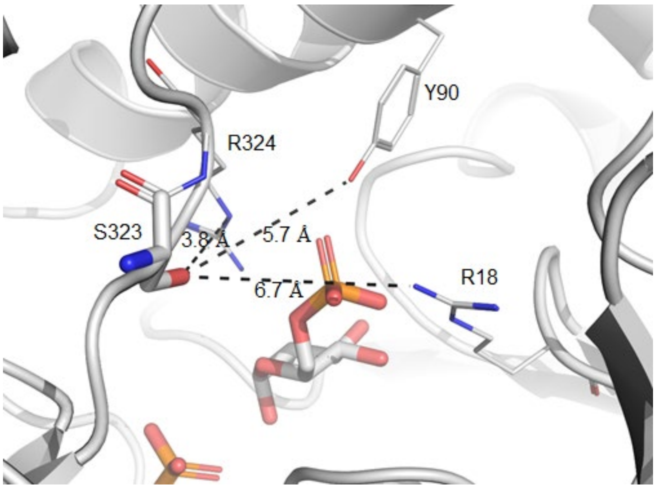


**Supplementary Figure S6:** View of the active site of *M. thermoresistibile* OtsA complexed with ADP and F6P (white). Distances between Ser323 hydroxyl group and other atoms interacting with the phosphate group of G6P are shown. Phosphorylation of Ser323 would place the phosphate group in a similar position to the one occupied by G6P.

**Table S2:** mCSM-lig predicitons

| Residues | Log change in affinity |
|---|---|
| ***V363F** | -1.4 |
| **D285E** | -1.3 |
| **T321P** | -1.3 |
| ***L319I** | -1.1 |
| ***E367L** | -0.8 |
| **P363H** | -0.6 |

*Residues selected for mutation. All residues except E367 are within 4.5 Å of the adenine moiety of ADP-glucose.

### Methods

**Table S3:** Primers used in this work

| | |
|---|---|
| *otsA*pET28SUMO_F | ATAGGATCCATGGCTGACCGGGGCGACTC |
| *otsA*pET28SUMO_R | ATTAAGCTTTCACACCGGAACCGCGTCGG |
| L319I_F | GACACCGTCTTCGTCCAGATCGCCACCCCCAGCCGCGAG |
| L319I_R | CTCGCGGCTGGGGGTGGCGATCTGGACGAAGACGGTGTC |
| V363F _F | CCTGCACCGGCCGTTTCCGCGTGAGGAAC |
| V363F _R | GTTCCTCACGCGGAAACGGCCGGTGCAGG |
| V363F_E367L_F | CCTGCACCGGCCGTTTCCGCGTGAGCTAC |
| V363F_E367L_R | GTAGCTCACGCGGAAACGGCCGGTGCAGG |
| R213E_F | CTTCCTGTTCCTGGCGCGAGAGCTGGTGGGCGCCAACAC |
| R213E_R | GTGTTGGCGCCCACCAGCTCTCGCGCCAGGAACAGGAAG |
| R384E_F | CATGCTGGTCACCCCGCTGGAGGACGGGATGAACCTGGT |
| R384E_R | ACCAGGTTCATCCCGTCCTCCAGCGGGGTGACCAGCATG |
